# Improving Peptide-Protein Docking with AlphaFold-Multimer using Forced Sampling

**DOI:** 10.1101/2021.11.16.468810

**Authors:** Isak Johansson-Åkhe, Björn Wallner

## Abstract

Protein interactions are key in vital biological process. In many cases, particularly often in regulation, this interaction is between a protein and a shorter peptide fragment. Such peptides are often part of larger disordered regions of other proteins. The flexible nature of peptides enable rapid, yet specific, regulation of important functions in the cell, such as the cell life-cycle. Because of this, understanding the molecular details of these interactions are crucial to understand and alter their function, and many specialized computational methods have been developed to study them.

The recent release of AlphaFold and AlphaFold-Multimer has caused a leap in accuracy for computational modeling of proteins. In this study, the ability of AlphaFold to predict which peptides and proteins interact as well as its accuracy in modeling the resulting interaction complexes are benchmarked against established methods in the fields of peptide-protein interaction prediction and modeling. We find that AlphaFold-Multimer consistently produces predicted interaction complexes with a median DockQ of 0.47 for all 112 complexes investigated. Additionally, it can be used to separate interacting from non-interacting pairs of peptides and proteins with ROC-AUC and PR-AUC of 0.78 and 0.61, respectively, best among the method benchmarked.

However, the most interestingly result is the possibility to improve AlphaFold by enabling dropout at inference to sample a wider part of the conformational space. This improves the median DockQ from 0.47 to 0.56 for rank 1 and the median best DockQ improves from 0.58 to 0.72. This scheme of generating more structures with AlphaFold should be generally useful for many application involving multiple states, flexible regions and disorder.

## 1 Introduction

Protein-protein interactions are central to all biological process, and understanding the molecular details of these interactions are crucial to understand and alter their function. Of protein-protein interactions, up to 40% are considered peptide-protein interactions, many of which are responsible for vital functions such as cell life-cycle regulation (Lee *et al*., 2019; Tu *et al*., 2015). These are interactions between a protein and a smaller peptide fragment, sometimes referred to as short linear motif or SLiM, which can be part of a disordered region of a larger protein. Because peptide fragments display a high degree of flexibility and are often disordered when unbound, investigating the molecular details of such interactions and indeed even identifying direct interaction at all has been difficult both from an experimental and computational point of view, and specialized solutions have been developed to investigate peptide-protein interactions specifically (Petsalaki and Russell, 2008; Alam *et al*., 2017; Lei *et al*., 2021; Öztürk *et al*., 2018; Johansson-Åkhe *et al*., 2021).

The unprecedented accuracy of AlphaFold (Jumper *et al*., 2021) and the recent release of the source code have transformed the field of computational and structural biology. It is now possible to achieve highly accurate protein structure prediction on par with experimental accuracy for many proteins, e.g. for human proteins 58% can be modeled accurately (Tunyasuvunakool *et al*., 2021), as a comparison the previously experimentally determined structures of human proteins covered 17%. This and the improved methods that will follow in the foot steps of AlphaFold will have a huge impact not only on structural biology but the whole life-science field.

Despite the fact that AlphaFold was trained on monomeric strucures, it has demonstrated an impressive ability and stability to allow manipulation of its input to predict protein complexes, i.e. using a 200 residue gap to infer a chain break (Bryant *et al*., 2021) or using a flexible linker (Ko and Lee, 2021; Tsaban *et al*., 2022). However, it is clear that even though the input-adapted versions of AlphaFold performed better than state-of-the-art, the recently released retrained AlphaFold-Multimer system (Evans *et al*., 2021) successfully predicts the interface (DockQ (Basu and Wallner, 2016) ≥ 0.23) for 67% of the cases. and at high accuracy (DockQ ≥ 0.8) for 23% of the cases on a set of 4,433 non-redundant interfaces, an improvement over the input-adapted version with 60% and 92%, respectively (Evans *et al*., 2021).

Although AlphaFold expertly implements several state-of-the-art neural network concepts and contributes their own with the Evoformer layers, the common bioinformatics concept of simple increased sampling of the conformational space with more lenient energy terms in search of a global optima is not explored. Repeated sampling of predictions with different random seeds or parameters is commonly utilized in bioinformatics methods, examples include Rosetta and ZDOCK (Raveh *et al*., 2011; Pierce *et al*., 2014). Activating dropout layers at inference has previously been suggested for introducing and mapping uncertainty while creating a model ensemble with no increase in training time (Gal and Ghahramani, 2016; Lakshminarayanan *et al*., 2017).

In this manuscript, we investigate whether AlphaFold’s milestones are applicable not only to the prediction of complexes of globular structures, but if AlphaFold and AlphaFold-multimer can also be used to further the field of peptide-protein interaction prediction as well as peptide-protein complex modeling. Additionally, we investigate the effects of increased sampling on the performance of AlphaFold, whether by increasing the recycling steps to allow AlphaFold to explore and refine the search-space by itself or by activating Dropout at inference to force variance in output.

## 2 Methods

### 2.1 Dataset

A test-set was constructed for the benchmarking of peptide-protein complex modeling which could also be used for peptide-protein interaction prediction.

The samples of observed peptide-protein interaction complexes in the test set were based on the state-of-the art paper for peptide-protein interaction prediction by Lei *et al*. (2021), which consisted of 262 peptide-protein complexes with experimentally solved structures in the PDB (Berman *et al*., 2000). After selecting one complex representative per ECOD (Schaeffer *et al*., 2017) family, the final redundancy reduced set consisted of 112 peptide-protein complexes.

For peptide-protein interaction prediction, a negative set was created in the same manner as in Lei *et al*. (2021); by randomly pairing 560 (so × 5 as many negatives as positives) peptides and protein receptors from the positive set. Although randomly pairing proteins is no guarantee for lack of interaction, the resulting false negative rate will be statistically insignificant, especially considering the proteins hail from different species, and is as such common practice for constructing negative sets for protein-protein interaction prediction (Wang *et al*., 2019; Guo *et al*., 2008).

This scheme results in a dataset which can both be used to evaluate the docking performance on the positive set as well as be used for benchmarking the capacity to predict whether peptide-protein pairs interact or not.

### 2.2 AlphaFold versions

One input-adapted version of AlphaFold monomer and several variants of AlphaFold-Multimer were included in the benchmark, as outlined below:

- *AF-gap*: This is an input-adapted version using AlphaFold (Jumper *et al*., 2021) monomer. The input is adapted by placing a 200 residue gap between the chains and pairing the MSAs diagonally (Bryant *et al*., 2021). The models were ranked by the average plDDT of the peptide as in Tsaban *et al*. (2022), and rank 1 was selected.
- *AFmulti-*: This is AlphaFold-Multimer version 2.1.0 (Evans *et al*., 2021). The models were ranked by the *ranking_confidence* score and rank 1 was selected. The *ranking_confidence* is a linear combination of the interface score, *ipTM*, and overall structural score, *pTM* : 0.8*ipTM* + 0.2*pTM*. AlphaFold multimer was run with one or several options from the list below:
  – *reduced_dbs*: using the reduced database setting.
  – *full_dbs*: using the large sequence databases, BFD and Uniclust30.
  – *template*: Also allowing templates as input to AlphaFold. Templates newer than 2019-10-14, E-values better than 10^*−*20^ to the target, or for the peptides *>* 95% sequence identity over the whole sequence, were filtered to ensure the target protein or close homologs could not be used.
  – *v2* : AlphaFold-Multimer version 2.2.0 (Evans *et al*., 2021, 2022) was run rather than 2.1.0.

So, if a variant of AlphaFold-Multimer is denoted as *AFmulti-v2_reduced_dbs_template*, this means that AlphaFold-Multimer version 2.2.0 was run with reduced database settings while allowing templates passing the template criteria to be used as input.

The difference between AlphaFold 2.1.0 and 2.2.0 is, according to the AlphaFold authors, that version 2.2.0 has been re-trained to produce less structural clashes while slightly improving performance. Additionally, 2.2.0 generates more structures by default (Evans *et al*., 2022). Since AlphaFold-Multimer version generates several structures per model by default, it was also run restricted to only producing one structure per model so as to enable a fair comparison to AlphaFold-Multimer 2.1.0. In this case, a *1* is added as a suffix to the variant name.

AlphaFold was run without the final relaxation step using Amber, thus only the unrelaxed models were evaluated.

#### 2.2.1 AlphaFold database versions

- Uniclust30 from `UniRef30_2021_06`
- Uniref90 from Aug 9, 2021.
- Uniprot, TrEMBL+SwissProt, from Nov 3, 2021.
- BFD database (Steinegger and Söding, 2018), clustering of 2.5 billion sequences from Uniprot/TrEMBL+SwissProt, Metaclust, Soil and Marine Eukaryotic Refence Catalog. Downloaded March 2019. `bfd_metaclust_clu_complete_id30_c90_final_seq.sorted_opt_cs219.ffindex MD5 hash: 26d48869efdb50d036e2fb9056a0ae9d`
- Mgnify, from `2018_12`.
- PDB, mmcif and seqres, from Nov 3, 2021 (restrictions where applied at run-time, see above).

### 2.3 InterPep2

InterPep2 (Johansson-Åkhe *et al*., 2020) is a template-based method for peptide-protein complex structure prediction, generating multiple peptide conformations and using TMalign (Zhang and Skolnick, 2005) and InterComp (Mirabello and Wallner, 2018) for finding structural templates of interaction and evaluating them by a random forest regressor. Models for the protein receptors were created using AlphaFold monomer and InterPep2 was run with standard settings but allowed no templates deposited into the PDB later than 2019-10-14, and additionally allowed no templates matching query protein receptors with an E-value better than 10^*−*20^ to ensure the query protein itself or particularly close homologs could not be used as templates for prediction. The random forest predicted suitability of the top-ranking template is used as the score for interaction prediction.

InterPep2 was run without the more computationally intensive refinement protocols, generating only coarse models. This will result in a lower number of sub-Ångström decoys but will only have minor to no effect on the positioning the peptide at the correct binding site (Johansson-Åkhe *et al*., 2020).

### 2.4 ZDOCK

ZDOCK is a fast and accurate FFT-based rigid-body docking method utilizing pairwise statistical potentials, and is a staple method for benchmarking protein-protein docking performance (Pierce and Weng, 2008; Pierce *et al*., 2014; Wallner and Mirabello, 2017; Moal *et al*., 2013; Yan and Huang, 2019). In this study, ZDOCK was used to generate 54000 decoys for each peptide conformation generated in the early steps of the InterPep2 protocol, a similar scheme as to that used in earlier ZDOCK studies and by ClusPro Peptidock or PIPER-FlexPepDock (Wallner and Mirabello, 2017; Porter *et al*., 2017; Alam *et al*., 2017), and the best-scoring decoy were selected as final prediction.

### 2.5 CAMP

The peptide-protein interaction prediction dataset of this study is constructed as in Lei *et al*. 2021 but redundancy-reduced, which presents a convolution neural network for peptide-protein interaction prediction called CAMP. As a benchmark reference in interaction prediction, the freely available model of CAMP run on all test targets. No retraining was performed, the model was run as is, as the test set is fetched from the same paper CAMP’s training set was defined in.

### 2.6 Performance Measures

#### 2.6.1 DockQ

To assess the quality of docked peptide-protein complexes, the DockQ program was used (Basu and Wallner, 2016). DockQ assesses the quality of a docked complex with regards to ligand root mean square deviation (LRMSD), the root mean square deviation of the interface residue conformation (iRMSD), and the fraction of native contacts recalled (fnat). The quality is measured on a scale of 0.0 to 1.0, with 0.25 representing models generally considered acceptably close to native, and 0.8 or above representing models with sub-Ångström quality, see Table 1. DockQ is not a linear measure; while extremely precise sub-Ångström changes in LRMSD can be required to increase DockQ-score from 0.7 to 0.8, requiring almost perfect structures to achieve higher DockQ-score, moving a completely flipped peptide towards roughly the correct binding site can be enough to raise DockQ-score from 0.1 to 0.2.

**Table 1:** Thresholds for DockQ measure of docked model quality.

| **DockQ** | **Model Quality** |
| --- | --- |
| $\geq 0.23$ | Acceptable |
| $\geq 0.50$ | Medium |
| $\geq 0.80$ | High |

#### 2.6.2 Receiving Operand Characteristic (ROC)

A ROC curve measures a method’s capacity to discover positive samples versus its number of misclassifications with a sliding score threshold, and plots true positive rate (TPR) versus false positive rate (FPR) for different threshold cutoffs. *TPR* = *Recall* = *TP/P* (*TP* =true positives, *P* =total positives in set). *FPR* = *FP/N* (*FP* =false positives, *N* =total negatives in set). The ROC area under the curve (AUC) is frequently used as a measure of classification method performance (Johansson-Åkhe *et al*., 2020; Lei *et al*., 2021).

#### 2.6.3 Precision, Recall, and F1

With an unbalanced dataset, TPR-FPR ROC curves and AUC can be used to benchmark methods against each other but fails to capture the actual absolute performance of the methods, for which Precision-Recall curves and AUCPR (AUC under such a curve, also referred to as *average precision*) should be used instead (Saito and Rehmsmeier, 2015; Davis and Goadrich, 2006). *Precision* = *TP/*(*TP* + *FP*). *Recall* = *TPR* = *TP/P*. F1 is the harmonic mean between precision and recall, defined as *F* 1 = (2*·Precision ·Recall*)*/*(*Precision*+*Recall*).

## 3 Results

In this study, the performance of AlphaFold when extended to the peptide-protein docking problem is benchmarked both comparing to previous established docking methods and comparing different ways to run AlphaFold. The benchmark was performed using a recently published data set of peptide-protein interactions (Lei *et al*., 2021) (see Methods for details). Multiple versions of AlphaFold including: *AF-gap, AFmulti-reduce_dbs, AFmulti-full_dbs, AFmulti-reduced_dbs_template, AFmulti-full_dbs_template, AFmulti-v2_reduced_dbs, AFmulti-v2_full_dbs, AFmulti-v2_full_dbs1, AFmulti-v2_reduced_dbs_template*, and *AFmulti-v2 full dbs template*; as well as InterPep2 (Johansson-Åkhe *et al*., 2020), and ZDOCK (Pierce *et al*., 2011) were included in the benchmark. All comparisons are performed on the targets for which all methods produced predictions, 112 complexes in total (see Methods for details).

### 3.1 AlphaFold is best with more data and sampling, but worse with templates

Model quality of the top ranked prediction from each method was assessed using DockQ (Basu and Wallner, 2016). The median DockQ is 0.47 for the best AlphaFold metod, AFmulti-v2 full dbs, compared to 0.12 and 0.08 for InterPep2 and ZDOCK, respectively, see Figure 1. AlphaFold-Multimer version 2.2.0 does produce more structures by default, sampling some nearby conformations, but AFmulti-v2 full dbs1 is still the highest performer of the remaining methods, albeit much closer to AlphaFold-Multimer version 2.1.0 with a median DockQ score of 0.42. AF-gap performs worse than AlphaFold-Multimer versions, with a median DockQ of 0.25, but that is expected since it was not optimized for this task.

**Figure 1:**
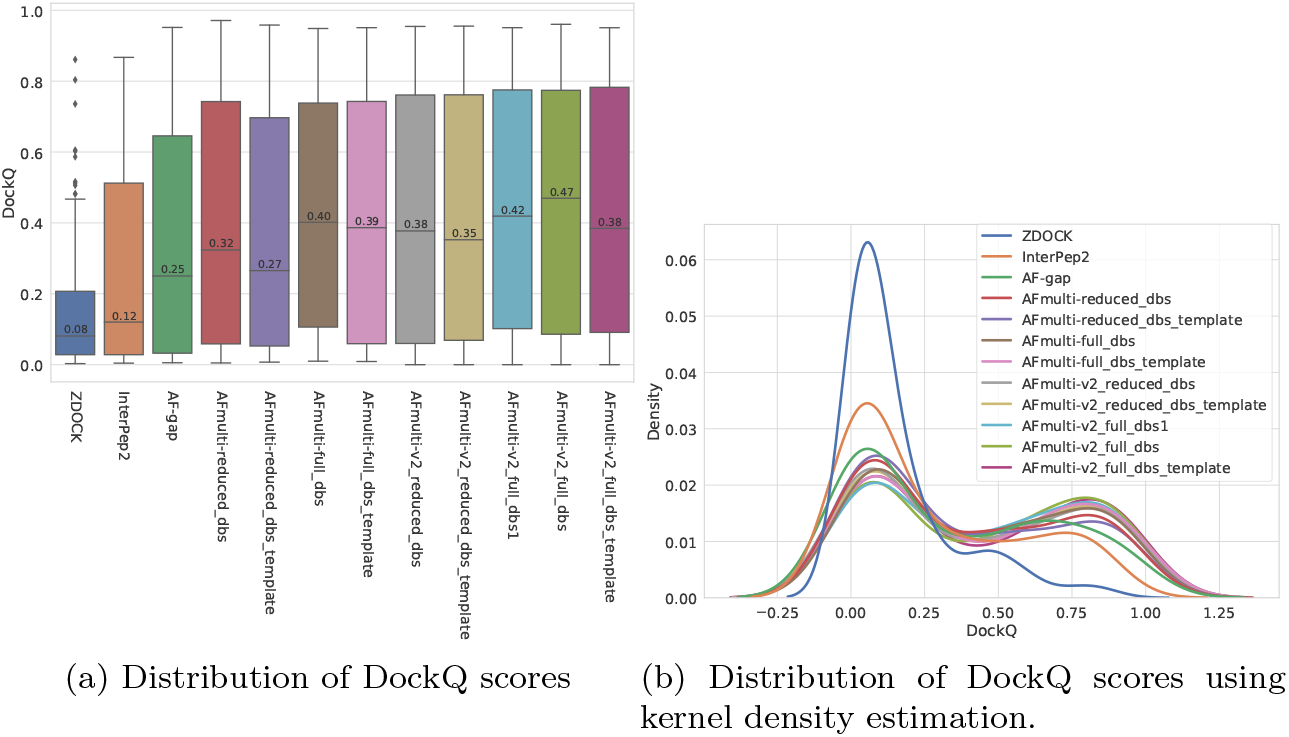
Model quality measured by DockQ

Of the different versions of AlphaFold-Multimer tested, it is consistently the versions with access to the larger databases (full dbs) that are superior. However, running AlphaFold without template information seems better than running it with template information, especially when also running with a reduced database size. Previous works have shown that although AlphaFold is fully capable of accurately judging the quality of structures derived from template information without any MSA information at all, it cannot efficiently sample the folding landscape without MSAs, and if given template information with no MSA information AlphaFold will essentially copy the template (Roney and Ovchinnikov, 2022). As the peptides are small in size, finding significant sequence matches is difficult, and their parts of the paired MSAs often have low number of effective sequences. This, in combination with the fact that the performance loss when including templates is greater when smaller databases are used for MSA construction, implies that the when run on peptide-protein complexes the protocol will become over-reliant on using templates to sample starting positions, and ought in such cases to be run without templates.

By examining the quality in more detail it can be seen that AlphaFold-Multimer version 2.2.0 (AFmulti-v2 full dbs) produces more Medium quality models compared to the other methods: 54 compared to only 43 and 30 for AF-gap, and InterPep2, respectively, see Figure 2, with minor advantages of other versions of AlphaFold-Multimer. Overall, the best AlphaFold-Multimer is able to predict at least an Acceptable model for 68/112 (61%), while the AlphaFold-Multimer with least information (AFmulti-reduced dbs) still produces 61/112 (54%) Acceptable models.

**Figure 2:**
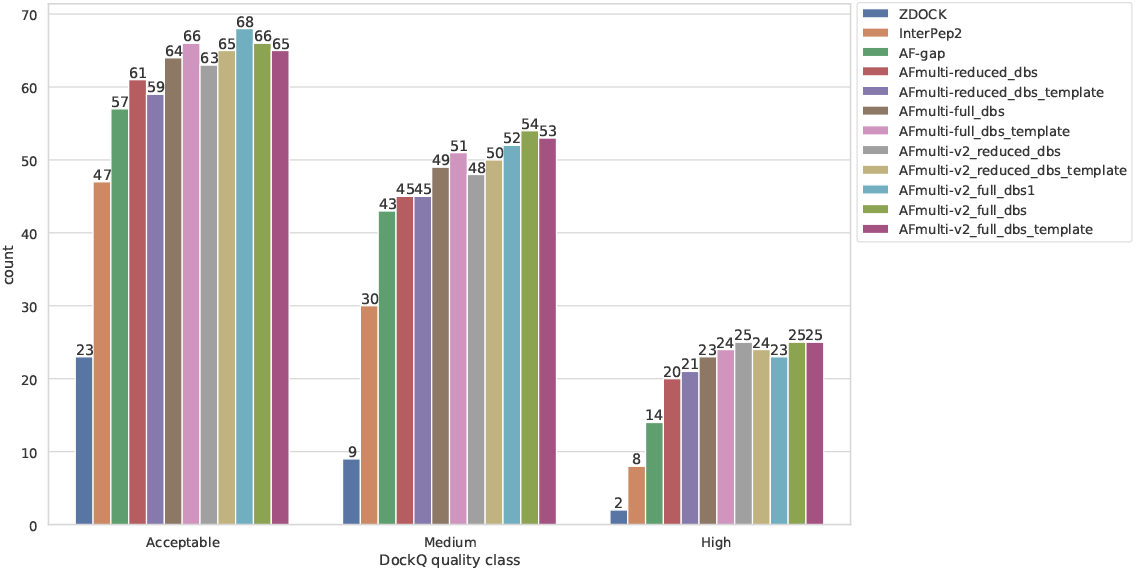
Distribution of model quality. For each DockQ-cutoff (Table 1), for how many of the complexes in the test set does the highest ranked model pass the cutoff threshold.

### 3.2 Forced Sampling Dramatically Improves AlphaFold

Two different methods were employed to force the best versions of AlphaFold-Multimer tested so far to sample more of the conformation space than the standard settings allow. First, the number of recycling steps run on the structure module layer was increased incrementally to explore AlphaFold’s own ability to improve its sampling by refining its predictions. Secondly, the dropout layers used during training of AlphaFold was turned on during inference and the network run several times in a Monte Carlo Dropout scheme, producing many structures for final comparison and evaluation. Such schemes have been used in the past to create ensemble methods from single trained models and has been used with classifiers to estimate model variance (Gal and Ghahramani, 2016).

As can be seen in Figure 3, both increasing the number of recycles and producing varied samples by running the network several times (nstruct>1) with dropout active have significant effects on performance, especially when combined. The improvement increases most rapidly for the the first 25 structures per network model (5×25=125 total), but for some settings there is a small but steady increase in DockQ all the way up to 200 structures (5×200=1000 total). The improvement is minor and we decided to stop at 200 structures in the interest of time, but it is certainly possible that even more structures could improve the method even further.

**Figure 3:**
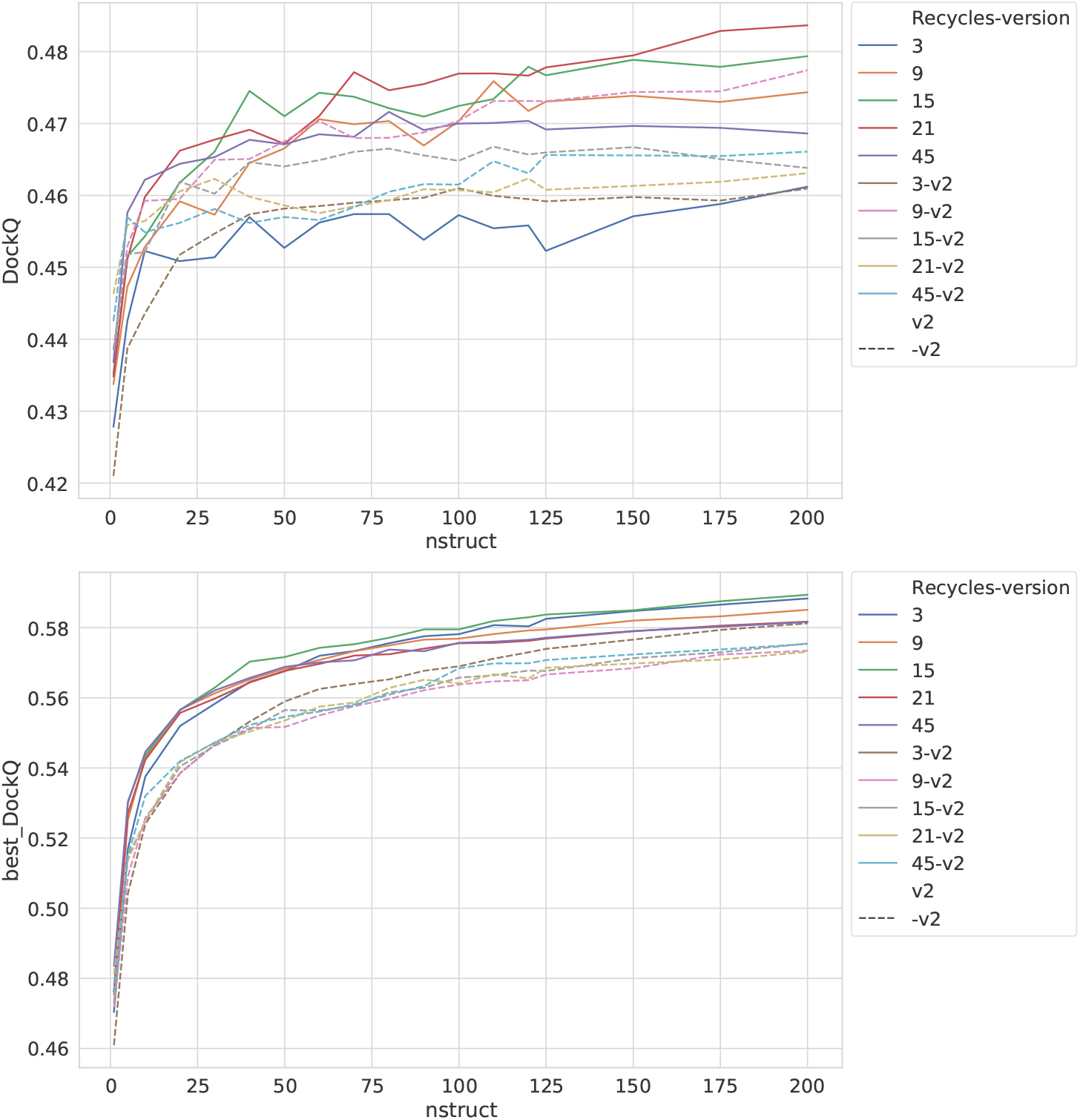
Effects on AlphaFold performance of increasing the number of recycles and allowing more sampling with dropout. The DockQ value reported in the left figure is the DockQ score of the model generated with the highest predicted score according to AlphaFold itself, and the value reported in the right figure is the best DockQ score of all models for each target. Note that the standard number of recycles for AlphaFold is 3, and running AlphaFold without dropout produces 5 models, since repeated runs offer little to no variance in the sampling and is unnecessary. The best AlphaFold without increased recycling or dropout scored an average DockQ of 0.42.

Dropout and increased recycles consistently leads to AlphaFold generating more varied models, including both more worse models but also better models than running just a single time without dropout. The difference between the best and worst DockQ is shown in Figure 4 for each target. In all cases there is a larger spread in favor of dropout. While dropout increase the sampling the correlation between the self-evaluating score and DockQ decreases somewhat from 0.74/0.71 to 0.72/0.67 for v1/v2, Figure 5. The correlation is still good and the better models can be picked out from the rest and overall performance increases, Figure 3.

**Figure 4:**
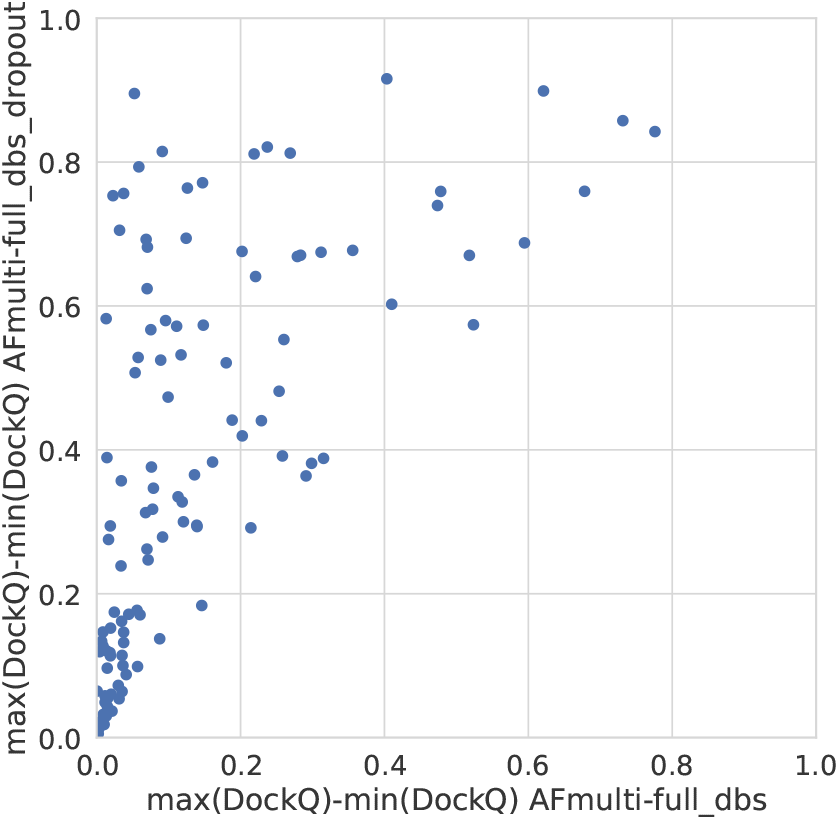
Difference between the maximum and minimum DockQ per target with and without dropout.

Additionally, combining dropout with increased recycles seems to restore the ranking performance in terms of correlation of AlphaFold-Multimer version 2.2.0 back up to correlation R of 0.71, while version 2.1.0 seems to simply generate more high-quality structures, Figure 5. Note, however, that there is still a substantial gap in median DockQ-score for selected models and best models generated, Figure 3. Indicating that AlphaFold is generating much better models than its scoring function is able to recognize and that there is potential to improve the method by improving the model quality assessment of the generated structures.

**Figure 5:**
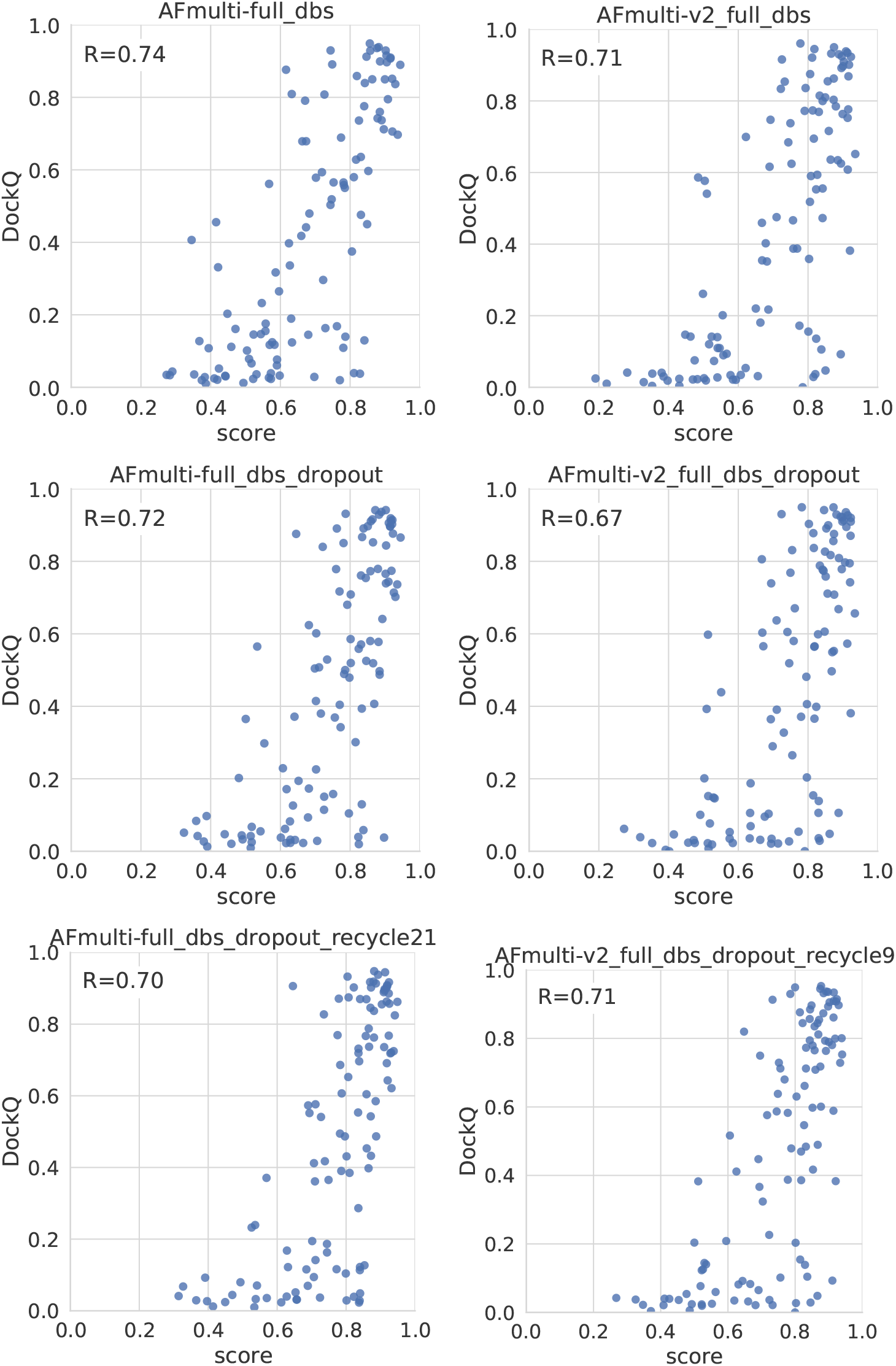
Correlation between method score and DockQ. The score is the *ranking confidence*.

From reminder of this study, whenever AlphaFold-Multimer is run with dropout, it produces 200 structures per neural network model (5×200=1000 structures in total). Whenever AlphaFold-Multimer-v1 is referenced as being run with additional recycles, it is run with 21 recycles unless otherwise specified. Similarly, AlphaFold-Multimer-v2 is run with 9 recycles. Note that the base behavior of AlphaFold is to run 3 recycles unless otherwise specified. These specific versions of AlphaFold-Multimer were selected based on their median DockQ score of their predictions, Figure 3.

### 3.3 Greater Improvements to AlphaFold by Selective Dropout

Even though running AlphaFold-Multimer with dropout active at inference improved the quality of final predicted docked peptide complexes, the correlation between AlphaFold predicted confidence and DockQ became worse (Figure 5). This is unsurprising, as dropout is not only applied to the evoformer layers of the network, but also the structure module responsible for translating the latent representation of the evoformer layers into the final protein structure as well as creating the final predicted confidence score. Roney and Ovchinnikov 2022 theorized that the purpose of the evoformer layers is to provide an initial low-energy guess of the protein structure, while the structure module refines this guess. If this theory holds, then including dropout at inference in the early layers of the AlphaFold network should be enough to introduce variance in final predictions and explore conformational space, while including dropout in the structure module might be used for uncertainty assessment but would have detrimental effects on final predicted complexes and in particular the score.

To test this, AlphaFold-Multimer versions 2.1.0 and 2.2.0 were run again on the entire dataset, using the best combination of hyper-parameters such as number of recycles and dropout seen in previous tests. This time, dropout was applied selectively to all parts of the network except the structure module, in hopes that the correlation between confidence score and DockQ would be retained in relation to running without dropout, thereby also improving the selection between generated structures and thus improving AlphaFold-Multimer for peptides overall. These versions are denoted with *dropout_noSM* for dropout except in the structure module.

Table 2 shows how for AlphaFold-Multimer version 2.1.0, this hypothesis holds true and correlation between predicted score and DockQ is restored while overall model quality is improved through selectively applying dropout rather than activating all dropout layers. With both versions 2.1.0 and 2.2.0, not applying dropout to the structure module seems to have no effect if the number of recycles is increased to optimal levels, indicating that predictive performance can be recovered with enough recycles.

**Table 2:** Correlations between predicted scores and DockQ of top predictions from different predictors.

| Predictor | Version | Dropout | Recycles | Corr. | Median DockQ |
| --- | --- | --- | --- | --- | --- |
| ZDOCK |  |  |  | 0.07 | 0.08 |
| InterPep2 |  |  |  | 0.76 | 0.12 |
| AF-gap | 2.0.0 | no | 3 | 0.42 | 0.25 |
| AlphaFold-Multimer | 2.1.0 | no | 3 | 0.74 | 0.40 |
|  | 2.1.0 | no | 21 | **0.75** | 0.43 |
|  | 2.1.0 | yes | 3 | 0.72 | 0.50 |
|  | 2.1.0 | yes | 21 | 0.70 | 0.54 |
|  | 2.1.0 | yes, noSM | 3 | **0.75** | 0.53 |
|  | 2.1.0 | yes, noSM | 21 | 0.70 | **0.55** |
| AlphaFold-Multimer | 2.2.0 | no | 3 | **0.71** | 0.47 |
|  | 2.2.0 | no | 9 | 0.68 | 0.49 |
|  | 2.2.0 | yes | 3 | 0.67 | 0.51 |
|  | 2.2.0 | yes | 9 | **0.71** | **0.53** |
|  | 2.2.0 | yes, noSM | 3 | 0.67 | 0.50 |
|  | 2.2.0 | yes, noSM | 9 | **0.71** | 0.50 |

The same table also shows that for AlphaFold-Multimer version 2.2.0, not including dropout in the structure module has a negative impact on final performance, and no impact on score correlation to DockQ. From the description of 2.2.0 from the authors of AlphaFold: “[the structures produced by this version] have greatly reduced number of clashes on average and are slightly more accurate.” (Evans *et al*., 2022). Minimizing clashes might thus be prioritized more strongly in version 2.2.0 over sampling a wider energy landscape, as opposed to version 2.1.0. As such, the dropout in the structural module might re-introduce some variance in the structures created and aid in the sampling of a wider landscape.

This can be interpreted similarly to a classic ramping energy function often used in Monte Carlo protein folding or docking protocols (Nivón *et al*., 2013; Raveh *et al*., 2011). In such protocols, it is common practice to use a more lenient or coarse-grained energy function, for instance by decreasing the weight of repulsive energy terms or a coarse-grained protein representation, during the early stages when a wider sampling of the conformational space is preferred. Later in these protocols, these terms are ramped up to negatively bias towards clashes and allow for more fine-grained refinement and production of more realistic models. With AlphaFold-Multimer version 2.1.0, performance is optimal if dropout is applied to earlier layers (yielding coarse-grained sampling of conformational landscape) but not in the structural module at the end (keeping a fine-grained energy function for the end of the protocol). With AlphaFold-Multimer version 2.2.0, the entire network is already more optimized towards fine-grained modeling, and as such dropout needs to be applied across the board to achieve varied enough sampling for the peptide-protein docking problem.

### 3.4 Complementary performance of AlphaFold Multimer v1 and v2

While both AlphaFold-Multimer versions 2.1.0 and 2.2.0 show higher performance with the increased sampling through dropout, with 2.1.0 seeing a larger improvement but 2.2.0 already starting at a higher performance level, the two versions show differences in their behavior overall. Version 2.1.0 performs better at high numbers of recycles and when the structure module is not subjected to dropout. Version 2.2.0 on the other hand seems to require dropout in all layers to perform optimally, and a generally lower number of recycles is better.

Since both versions produce predictions in the same predicted score range and both correlate their scores well with DockQ, it is trivial to construct combination predictors by allowing the different versions to generate 100 structures each and then selecting the structure with the overall highest predicted score (200 structures total, for fair comparison to other versions). Different such combinations can be compared to the versions of AlphaFold-Multimer investigated so far in Figure 6. Indeed, such a simple combination manages to significantly increase performance even further than the increased sampling alone, raising it from median DockQ scores of 0.531 or 0.549 to 0.562 for the best combination. The best combination consists of the versions with best correlations between predicted score and DockQ score, AFmulti-full dbs dropout noSM and AFmulti-v2 full dbs dropout recycle9, and even though AFmulti-full dbs dropout noSM recycle21 is the version of AlphaFold-Multimer 2.1.0 with best median DockQ it is not part of the best combination. At this point, it seems as though correlation between ranking score and DockQ is more important than median model quality of models generated, as models are already generated with far higher quality than what is selected (Figure 7b).

**Figure 6:**
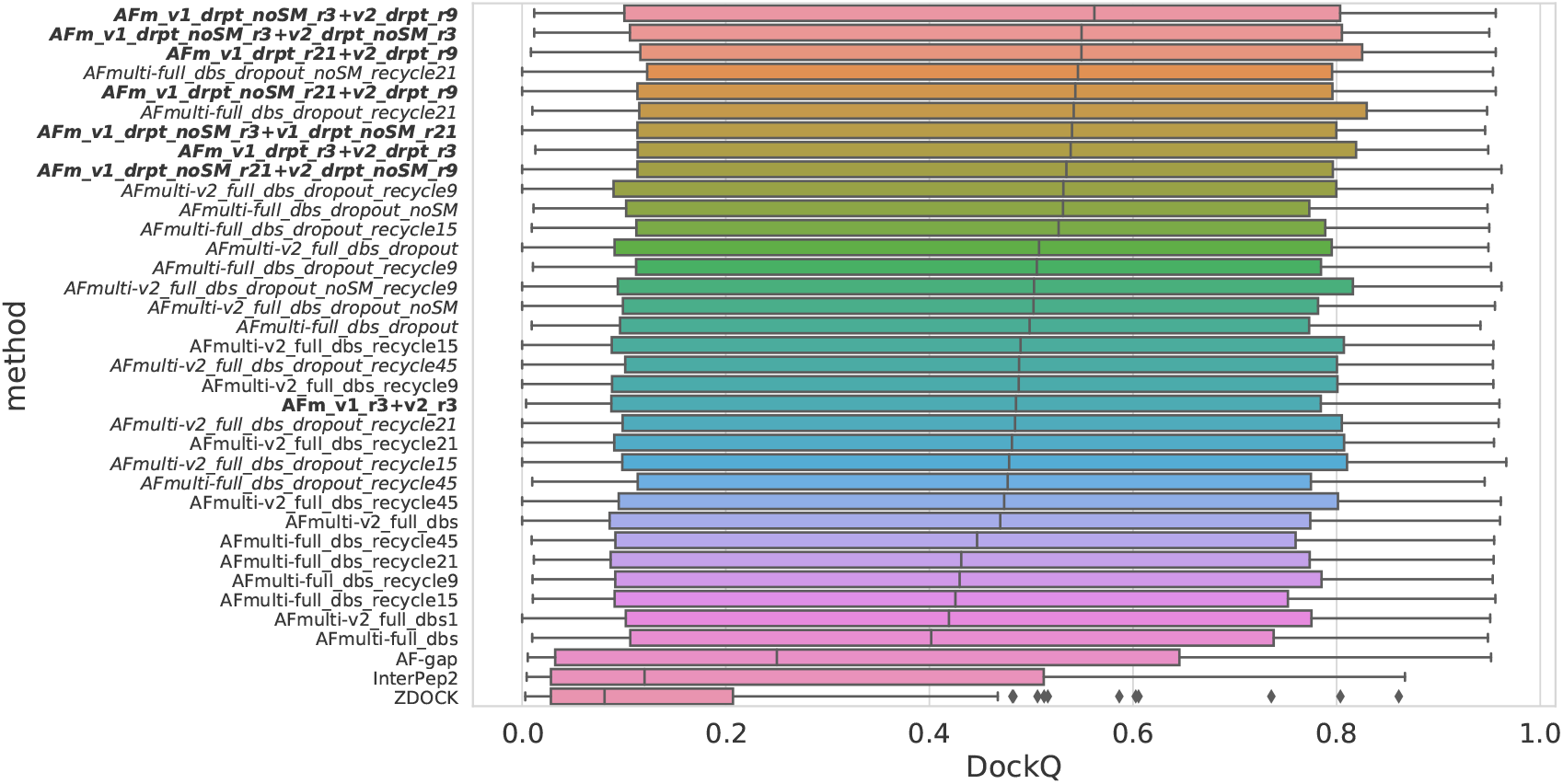
Boxplot of distributions of DockQ scores for final selected predictions for each target. Variations on AlphaFold-Multimer version 2.2.0 are marked in italics while combinations of versions 2.1.0 and 2.2.0 are marked in bold. The contractions *drpt* and *r* stand for *dropout* and *recycles*, respectively.

**Figure 7:**
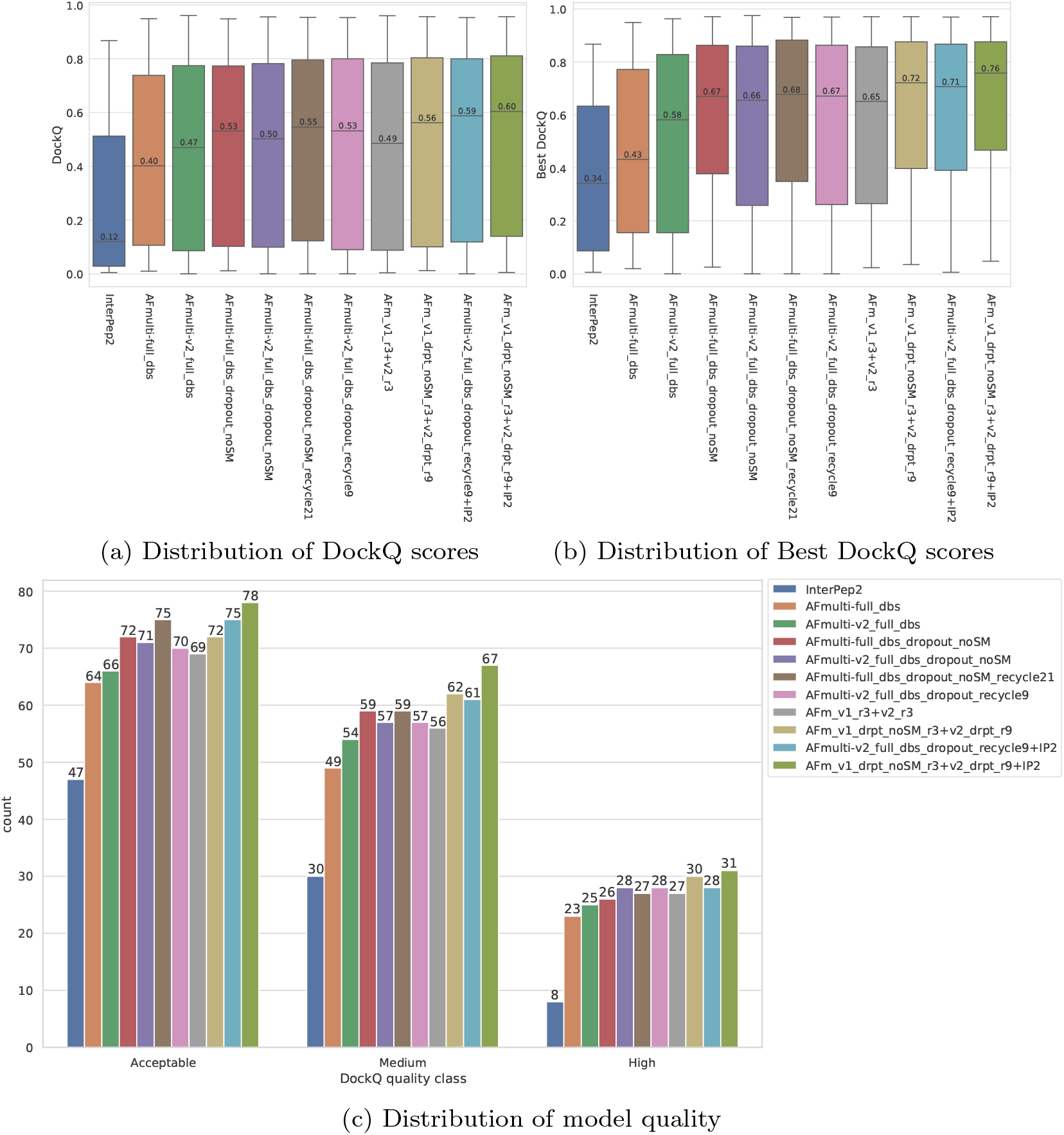
Improvements to AlphaFold-Multimer prediction of peptide-protein docking while applying the simple improvements described in this paper are significant.

#### 3.4.1 Complementary performance of AlphaFold Multimer and InterPep2

While the performance of AlphaFold-Multimer, especially with the improvements described in this study, is far above the template-based InterPep2, both AlphaFold-Multimer and InterPep2 possess scores which correlate well with DockQ for their predictions. A simple meta-predictor can easily be designed by assigning a cutoff to AlphaFold-Multimer’s predicted score after which InterPep2 will be run instead. Similarly, if that method also indicates a prediction failure according to its own cutoff, the safest choice is to fall back on the original AlphaFold predictions, as they are more likely to have higher DockQ. Cutoffs were selected by leave-one-out cross-validation, which surprisingly put them at approximately the suggested cutoffs for the individual papers for AlphaFold and InterPep2: 0.7 for AlphaFold and 0.4 for InterPep2.

The results of this improved meta-predictor for peptide-protein docking can be found in Figure 7. The performance increase from including InterPep2 is surprisingly substantial, raising the median DockQ from 0.562 to 0.604 even though InterPep2 by itself produces models with a way lower median DockQ than all AlphaFold versions tested. This probably stems from the high correlation between AlphaFold and InterPep2 predicted scores and DockQ score. Indeed, when looking at the best models sampled for the best combination methods, they all reach median DockQ scores in excess of 0.7. The final selected models, however, have median DockQ scores of only up to 0.6 in the case of the very best combination. Conformations of higher quality are obviously sampled, but not selected by the predicted score, indicating yet again that for running AlphaFold with dropout, the remaining challenge is more of a model ranking or quality assessment problem than a structure prediction problem.

The final meta-predictor should prove extremely useful to the peptide-protein docking field, as even AlphaFold-Monomer could, when adapted for peptide-protein docking through use of a poly-glycine linker, show similar performance to state-of-the-art PIPER-FlexPepDock (Tsaban *et al*., 2022).

### 3.5 AlphaFold can be used for Interaction prediction

Since the AlphaFold-Multimer predicted score correlates well with the DockQ score of the predicted complexes and the median DockQ of predicted complexes for true interactions is well above the Acceptable DockQ cutoff, it might be possible to use AlphaFold as an interaction predictor.

The positive set used to assess model quality above was merged with the negative set containing peptide-protein pairs that do not bind (see Methods). The scores for binding vs. non-binding pairs are shown in Figure 8. In general, pairs that bind are scored higher than the negative set across all methods. The Precision-Recall-AUC curve is 0.640 for AFm v1 r3+ v2 r3 median+InterPep2 compared to 0.615 for AFm_v1_r3+_v2_r3_median and 0.608 for the best single AlphaFold-Multimer version (AFmulti-v2_full_dbs_median), showing that here too, the combination of the methods improves performance. Additionally, basing the interaction prediction on an ensemble by picking the median predicted score rather than the highest predicted score seems to yield slightly better performance, Figure 9.

**Figure 8:**
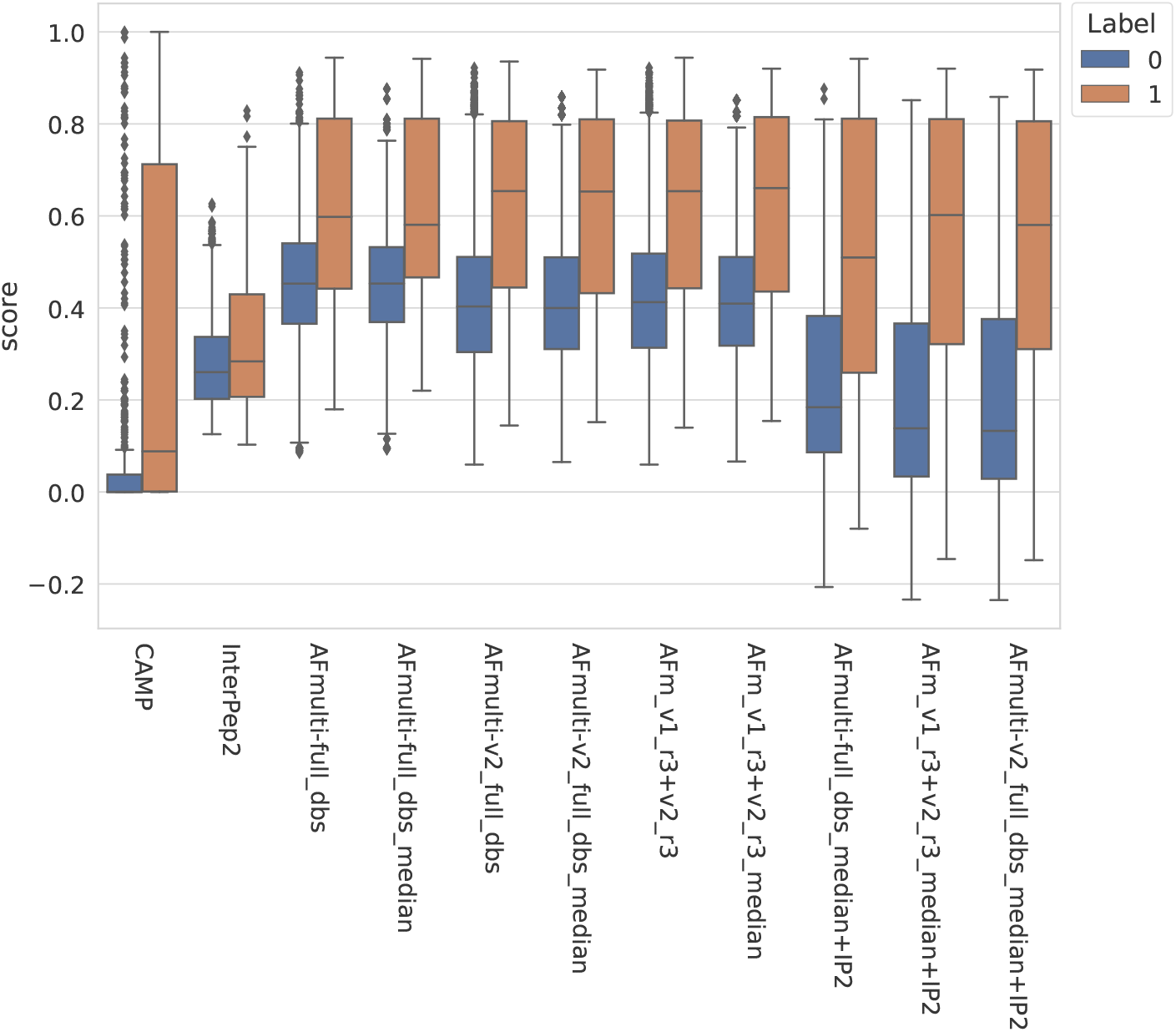
Score distribution for binding (label: 1)/non-binding (label:0)

**Figure 9:**
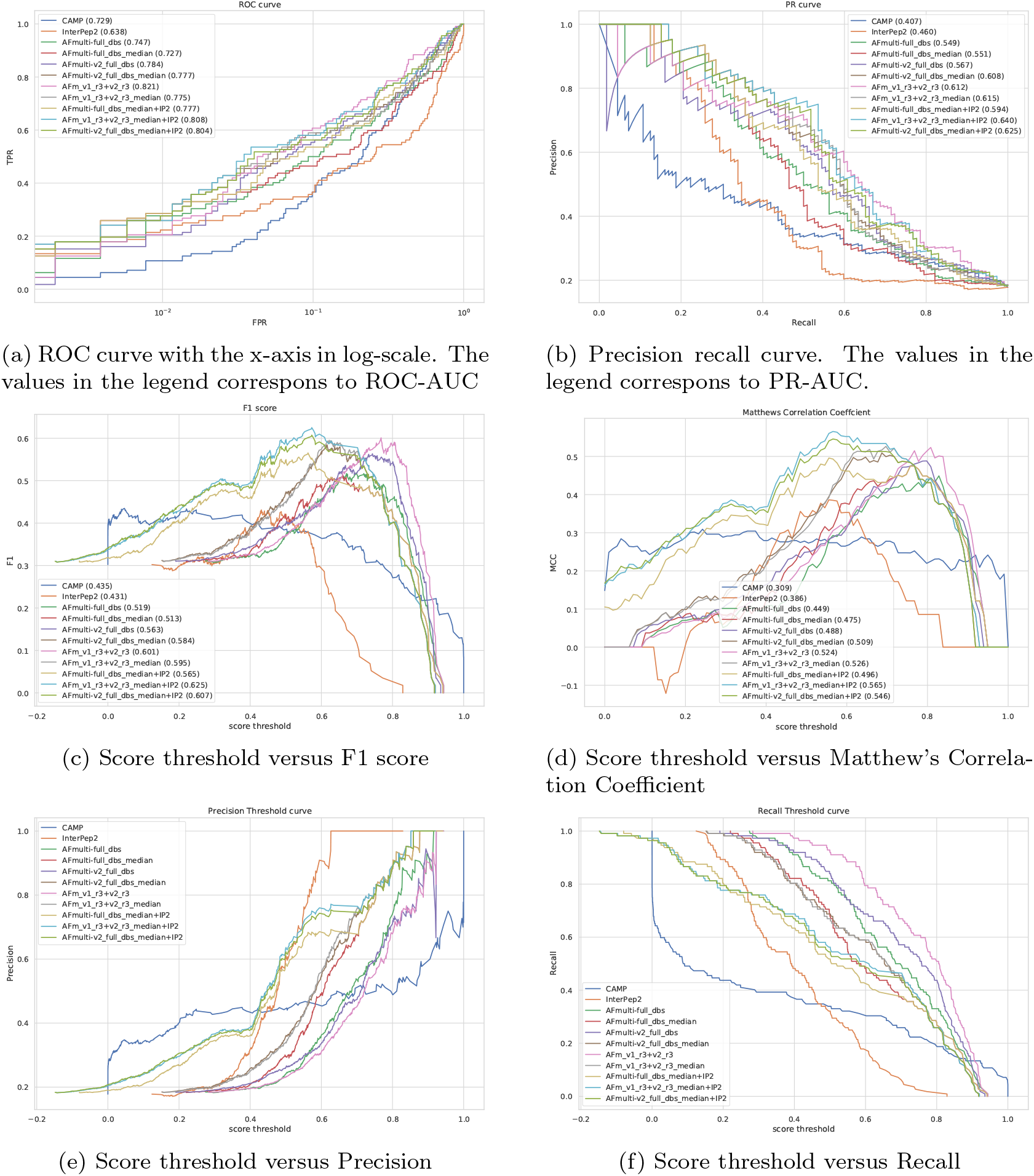
ROC curve, PR-curve, and plots showing score thresholds versus different metrics.

Performance for interaction prediction needs to be assessed at low FPR to enable all-vs-all comparisons with high confidence predictions without too many false positives. At FPR=0.01 the best methods recall around 0.3(TPR) of the positive examples, see Figure 9. From the Precision-Recall curve 0.3 recall corresponds to 0.83 precision for the best methods; and the corresponding to the score thresholds for the methods are 0.78, 0.50, and 0.78 for AlphaFold-Multimer (v1 and v2), InterPep2, and AFm_v1_r3+_v2_r3_median+InterPep2, respectively, see Figure 9. At the score threshold for low confidence prediction in AlphaFold (0.70) the precision is around 0.7 if the median score of the generated structures is used if instead the first ranked score is used the precision drops to 0.47, meaning that a prediction over that threshold has close to a fifty-fifty chance of being correctly predicted as interacting. The low confidence threshold for InterPep2 is 0.4 and corresponds to precision 0.3.

### 3.6 Example - Improvements for AlphaFold

To understand why AlphaFold fails at certain samples as well as why forced sampling through dropout and the template-based method InterPep2 can help with prediction quality in these cases, cases where AlphaFold failed were examined. One such example can be seen in Figure 10a, where AlphaFold has positioned part of the peptide correctly, but has flipped the orientation of the peptide and positioned the rest of it outside its binding pocket, as if to continue the peptide chain that direction. In fact, in 50% of cases where the AFmulti-v2 full dbs dropout recycle9+IP2 method produced an improved model of at least acceptable quality while AFmulti-v2 full dbs dropout recycle9 alone failed, the AlphaFold model positioned the peptide close to or at the correct binding site but in the wrong direction.

**Figure 10:**
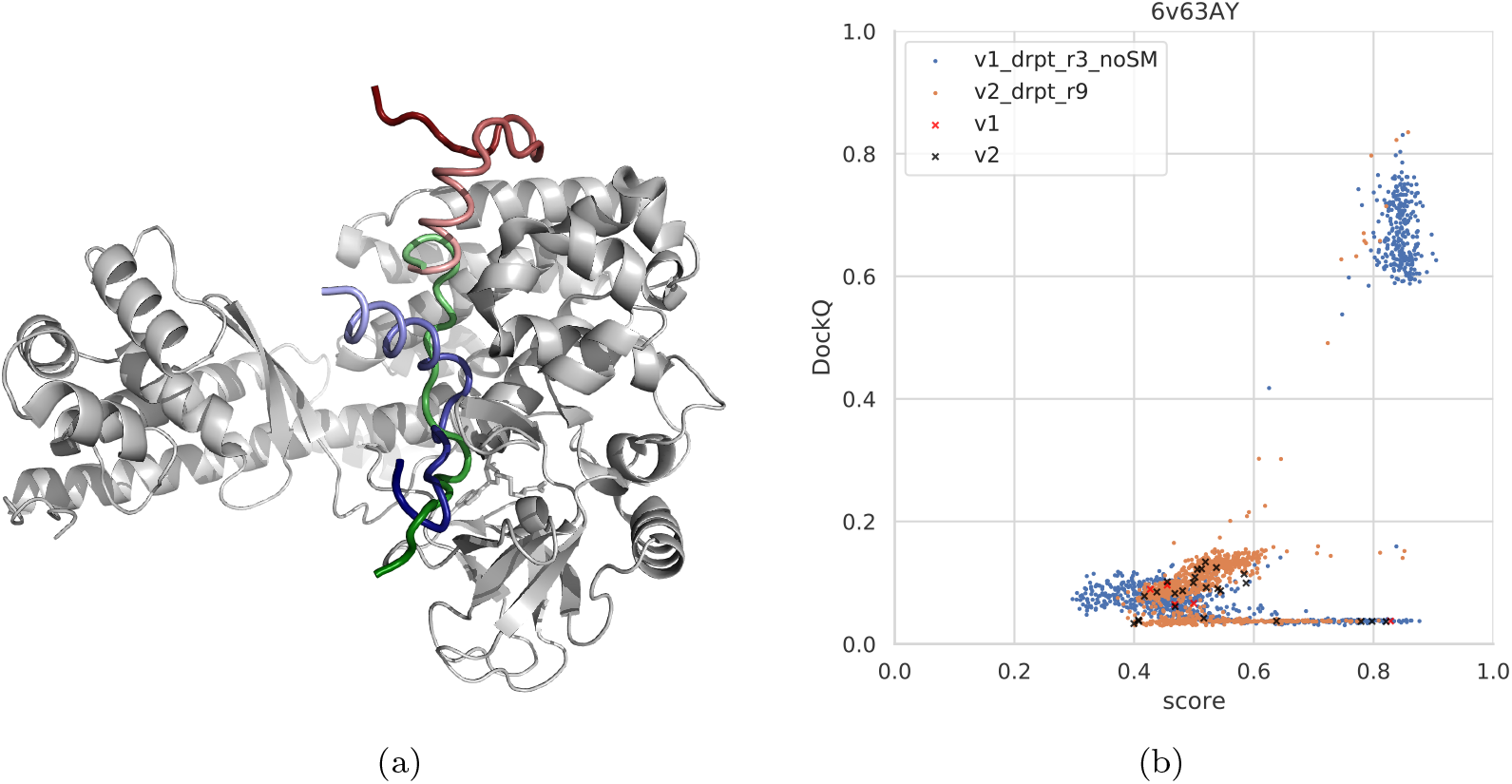
**(a)** An example from the test set where both forced sampling through dropout and inclusion of InterPep2 as a simple meta-predictor can improve the prediction. The predictions are of a cytoplasmic actin peptide binding to actin-histidine N-methyltranferase (PDB ID 6v63). The top predictions by AlphaFold-Multimer positions the very end of the C-terminal in the correct site (top1 prediction in red and native peptide colored green), but positions the peptide in the opposite direction of the actual fold, leading to most of it binding outside the pocket. The prediction by InterPep2 (blue) is coarse with some minor clashes, but has positioned the peptide inside the correct binding site. Note how both AlphaFold and InterPep2 have adopted similar folds for the peptide. **(b)** Scatterplot of AlphaFold-Multimer predicted score versus DockQ-score for all models generated for the 6v63 complex. When dropout at inference forces increased sampling, some conformations are sampled in the correct binding pose.

In this particular case, increased sampling through forced dropout with AlphaFold-Multimer version 2.1.0 could also result in the correct conformation being sampled and selected, as AlphaFold samples more nearby conformational space including the reverse direction of the peptide (and some states half-way rotated). In fact, the increased sampling for version 2.2.0 also samples a few high-DockQ conformations, although none of them are scored high enough to be selected (Figure 10b).

In 33% of improved cases, AlphaFold positioned the peptide at the binding site of a non-peptide binding partner, either for a co-factor or another protein chain (dimerization site, for instance). AlphaFold seems able to model the receptor protein regardless of the presence of any other binding chains or co-factors (Jumper *et al*., 2021; Tunyasuvunakool *et al*., 2021). However, as is evident from these examples, the absence of other binding partners and co-factors from the AlphaFold-Multimer modeling seems to result in a negative bias on the docking performance. Co-factors and other binders being absent from a complex seems to have less effect on the performance of the purely template-based method InterPep2, perhaps explaining in part why it is a good complement to AlphaFold-Multimer.

## 4 Conclusions

We have shown that AlphaFold-Multimer achieves state-of-the-art performance in both peptide-protein docking and peptide-protein interaction prediction without modification. Most interesting however, is the discovery that forcing increased sampling of the folding space through increasing the number of recycles or adding dropout at inference and running the protocol several times can improve performance of AlphaFold significantly. For the peptide-protein docking problem, with many degrees of freedom for the peptide and sometimes dynamic binding modes, the improvement in median DockQ score is dramatic, from 0.40/0.42 to 0.55/0.53 for version v1 and v2, respectively. We also demonstrated that it is possible to combine AlphaFold with InterPep2 to improve the median DockQ to 0.60. These results reinforce the previous findings that AlphaFold is well-suited for the peptide-protein docking problem which require a wider sampling of conformations. However, the improved sampling protocol presented here is not limited to peptide-protein docking problem and should be useful in many AlphaFold applications, to investigate multiple stable conforma-tions, for larger and more difficult targets, or for larger assemblies. Additionally, more variations in the application of dropout at inference could be investigated, such as different rates of dropout.

While the improvements presented here are significant, there is still a large gap between the quality of the best model generated and the one ranked highest by AlphaFold’s predicted score. As such, protein model quality assessment remains an important field of research in the future of protein structure prediction with AlphaFold.

## Acknowledgments

This work was supported by a Swedish Research Council grant, 2016-05369 and 2020-03352, The Swedish e-Science Research Center, and Carl Tryggers stiftelse för Vetenskaplig Forskning, 20:453. The computations were performed on resources provided by the Swedish National Infrastructure for Computing (SNIC), Knut and Alice Wallenberg (KAW), and Linköping University (LiU) at the National Supercomputer Center (NSC) in Linköping.

